# Molecular mechanism of Aurora-A binding and activation by hnRNPK

**DOI:** 10.64898/2026.10.11.757697

**Authors:** Isha A. Mohan, Lily Martin Viney, Matthew Batchelor, Mark W. Richards, Takashi Ochi, Richard Bayliss

## Abstract

Aurora-A kinase (AurA) is a serine/threonine kinase regulated by protein binding partners such as TPX2, which interacts with the Y-pocket of the kinase domain. Although the canonical functions of AurA are mitotic, where it phosphorylates spindle-associated proteins such as TACC3, there is evidence for non-mitotic, chromatin-associated roles. Heterogenous nuclear ribonucleoprotein K (hnRNPK) is a transcriptional regulator and an AurA substrate, how AurA is activated in this pathway is unknown. Their interaction contributes to cancer-associated nuclear accumulation of AurA and has been implicated in driving oncogenic transcription of MYC. Here, we show that hnRNPK promotes AurA autophosphorylation and stimulates AurA activity. The interface was mapped to residues 330-380, an intrinsically disordered region of hnRNPK, and the AurA Y-pocket, using NMR spectroscopy and competitive-binding experiments with mitotic and synthetic AurA-binding partners. AlphaFold3 modelling of the complex predicted Phe339 and Trp345 at the kinase Y-pocket, and site-specific mutagenesis confirmed these as key residues for AurA binding and activation. Our work provides a mechanistic basis for hnRNPK-dependent activation of AurA and efficient phosphorylation of hnRNPK Ser379. Defining the AurA/hnRNPK interface highlights the Y-pocket as a target site for modulation of nuclear AurA, a potentially cancer-specific role of the kinase.

**Significance Statement:** Aurora-A kinase (AurA) has critical functions during mitosis but is increasingly connected with cancer-associated roles in the cell nucleus. One such pathway involves the transcriptional regulator hnRNPK, which functions with AurA to promote oncogene expression, yet how the kinase is activated in this context has remained unclear. We show that an intrinsically disordered region of hnRNPK directly activates AurA through interaction with a regulatory site in the kinase domain. Through defining the molecular basis of this interaction and identifying mutations that disrupt it, we provide a mechanistic explanation for AurA activation in the cell nucleus and establish a framework for understanding how AurA contributes to oncogenic transcription.

## Introduction

Certain oncogenic functions of Aurora A kinase (AurA) can be attributed to the well-described and physiologically essential aspects of mitotic regulation carried out by pools of AurA in the cytoplasm, at the centrosome and on the spindle apparatus. The presence of AurA in the cell nucleus has been recognised as a hallmark of some cancers and a histological marker predicting poor survival (Aradottir et al., 2014). Indeed, in a meta-analysis, 8 of 18 tumour types were reported to have prominent nuclear AurA (Naso et al., 2021). Likewise, exclusion of AurA from the nucleus by fusion of a nuclear export signal has been shown to abolish its oncogenic activity (Tatsuka et al., 2009). These observations suggest that key roles of dysregulated AurA in cancer are carried out in the cell nucleus, thus selective targeting of its nuclear interactions could provide new avenues for development of more selective, AurA-directed cancer therapeutics.

Aberrant expression of hnRNPK is associated with progression and poor prognosis in cancer and its depletion decreases tumour growth (Tang et al., 2014). Its multiple cellular roles include transcriptional regulation through direct binding to the genome, particularly at cytosine-rich CT-elements, and in cancer it has been implicated in driving expression of oncogenes such as *MYC* and *c-src* (Michelotti et al., 1996; Wang et al., 2020). *MYC*, which encodes the key oncogenic transcription factor protein c-Myc, is driven by two promoters, P1 and P2, with a preference for P2 in normal cells (Levens, 2008). Binding of hnRNPK to CT-elements associated with the P1 promoter results in a shift in promoter usage and enhanced *MYC* transcription. Using Chromatin immunoprecipitation (ChIP) assays, Zheng et al. (2016) identified the presence of AurA at these sites and confirmed that AurA is recruited there through interaction with hnRNPK and is required for hnRNPK-mediated transactivation of the *MYC* P1 promoter. AurA had previously been shown to directly interact with hnRNPK and phosphorylate it at Ser379, which sits within an RGSY motif that is consistent with the consensus AurA substrate sequence R-X-S/T-ϕ, (where X is any residue and ϕ is any hydrophobic residue except proline) (Alexander et al., 2011). Thus, a stable nuclear complex of AurA kinase and hnRNPK has been shown to form in cancer cells that drives oncogenic *MYC* expression and otherwise influences the transcriptional programme to promote a cancer stem cell phenotype (Zheng et al., 2016). The mechanistic basis of the interaction between AurA and hnRNPK remains unresolved. In the present work, we show that hnRNPK is an activator of AurA kinase activity, map the molecular determinants of AurA and hnRNPK that mediate their interaction, and present a plausible molecular model of the complex, supported by experimental data, that provides a starting-point for its therapeutic targeting.

## Results

hnRNPK is a 463 amino acid protein that contains three nucleic acid-binding K homology (KH) domains and disordered regions incorporating a nuclear localisation signal (NLS) and a proline-rich K interacting (KI) domain (Figure 1A). To confirm that AurA and hnRNPK interact directly, we carried out co-precipitation experiments using a biotinylated Avi-tagged AurA kinase domain protein incubated with full-length hnRNPK^1-463^ (Figure S1A). A C-terminal fragment, hnRNPK^330-463^, which begins near the end of the KI region and includes the KH3 domain, was shown to retain AurA-binding activity using size-exclusion chromatography (Figure 1B). ^15^N,^13^C-labelled hnRNPK^330-463^ was expressed and purified, and chemical shift assignments were made using standard ^1^H-based nuclear magnetic resonance (NMR) spectroscopy experiments. Analysis of secondary structure using measured chemical shifts showed the expected three α-helix, three β -strand structure of the folded KH3 domain (Figure 1C, see PDB: 1KHM). The N-terminal part of the fragment, corresponding to residues 330-385, was disordered except for residues 355-360, which exhibited some weak α-helical propensity (Figure 1C). Titration of AurA protein into hnRNPK^330-463^ induced peak intensity reductions and chemical shift perturbations (CSPs) in ^1^H–^15^N-HSQC (or ^1^H–^15^N-TROSY) spectra (Figure 1D-F). Clear CSPs were observed for Trp345, Asp346 and a few other residues towards the N-terminus of the disordered region (Figure 1D, E). Corresponding peak intensity losses were strongest for hnRNPK residues 342-361, and to a lesser degree 369-374 (Figure 1F). Significant peak intensity losses were observed upon addition of only 0.1 mole fraction of AurA, indicating a dynamic mode of interaction. No CSPs and less significant peak intensity losses were observed for residues 385-463, indicating that the KH3 domain is not involved in AurA binding, consistent with the behaviour of this fragment in co-precipitation experiments (Figure S1B).

**Figure 1.**
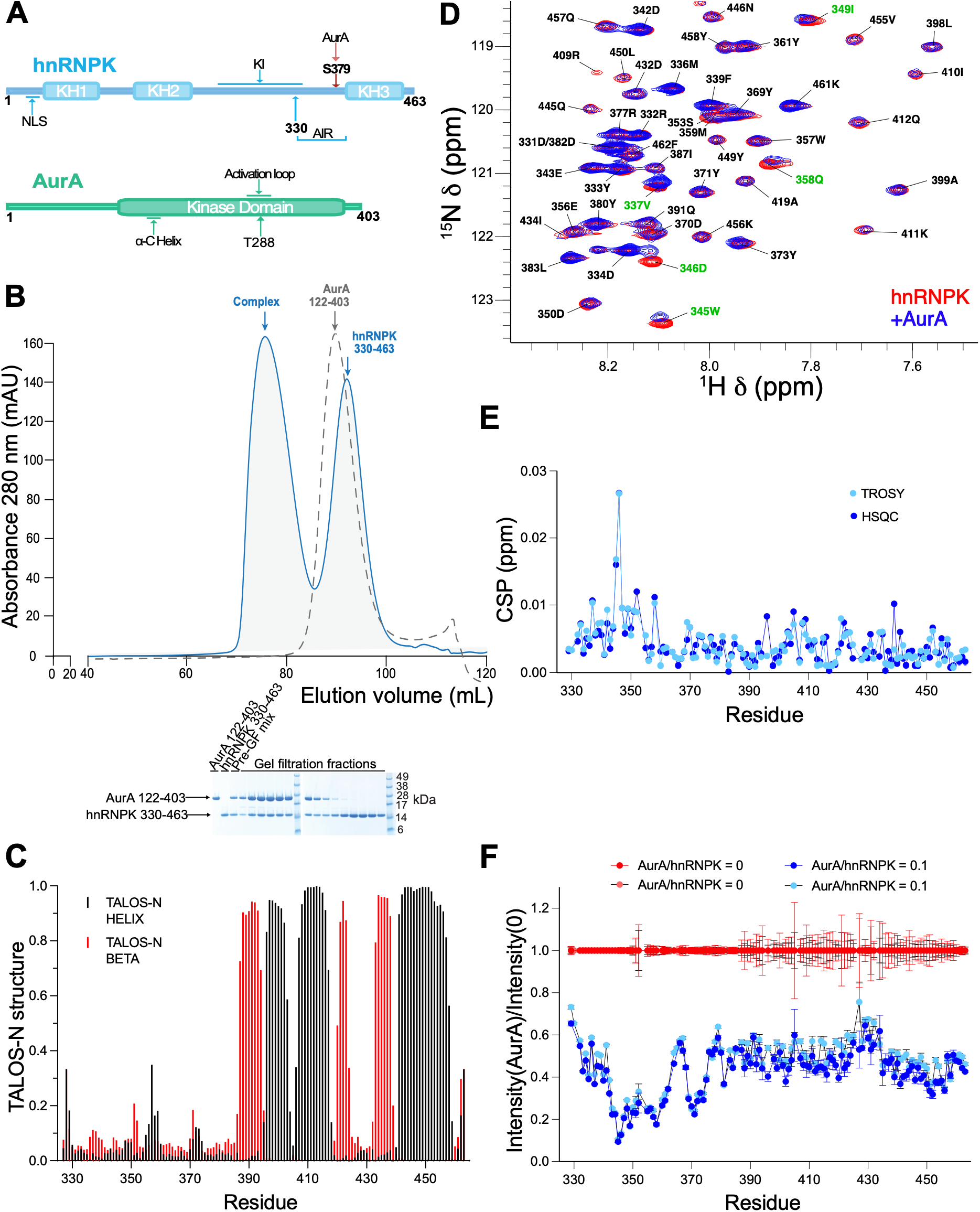
Biochemical and NMR characterisation of the interaction between AurA and hnRNPK. **(A)** Schematic showing the domain structures of hnRNPK, with the identified AurA interaction region (AIR), and AurA. (**B)** Size-exclusion chromatography (SEC) purification of an AurA/hnRNPK complex. Superposed chromatograms of AurA alone (grey dashed) and AurA/hnRNPK (blue). Excess hnRNPK is excluded as an hnRNPK-only peak. **(C)** TALOS-N NMR structural analysis of hnRNPK 330-463 using measured chemical shifts and sequence information to estimate the fractions of α-helix (black) and β -strand content (red) across the sequence. **(D-F)** NMR data for hnRNPK 330-463 before (red) and after addition of 0.1 mole fraction AurA (blue). **(D)** Part of the ^1^H-^15^N HSQC spectrum **(E)** Chemical shift perturbations (CSPs) and **(F)** Peak intensity ratios for hnRNPK residues upon addition of AurA. Results for HSQC and TROSY experiments are colour-coded red/dark blue and pink/light blue, respectively.

Having determined that the primary AurA-interacting region (AIR) of hnRNPK comprises residues within the 330-380 disordered region, a structural model of the complex (Figure 2A) formed by these proteins was generated by supplying AlphaFold3 (Abramson et al., 2024) with one copy of the sequence of AurA^122-403^, one copy of hnRNPK^330-380^, ADP and two Mg^2+^ ions. In the largest structurally-consistent set of models, which accounted for more than a quarter of the 100 models generated, the N-terminal portion of hnRNPK^330-380^ is predicted to bind AurA mainly through insertion of the side chains of Phe339 and Trp345 into a pocket on the N-lobe previously identified as the Y-pocket, part of the binding site of the AurA-activator Targeting protein for XKlp2 (TPX2) (Bayliss et al., 2003) (Figure 2B). Within the C-terminal portion of hnRNPK^330-380^, which is not consistently or confidently predicted to be involved in binding at the surface of AurA, residues 354-361 are modelled as an α-helix, consistent with NMR data that identified localised α-helical propensity in this region (Figure 1C). Also consistent with NMR data, the region C-terminal to this α-helix, incorporating the Ser379 site that is phosphorylated by AurA, is predicted to be disordered. This unbound region is sufficiently long and flexible that Ser379 could reach the active site of the same AurA protomer in order to be phosphorylated (Figure 2A).

**Figure 2.**
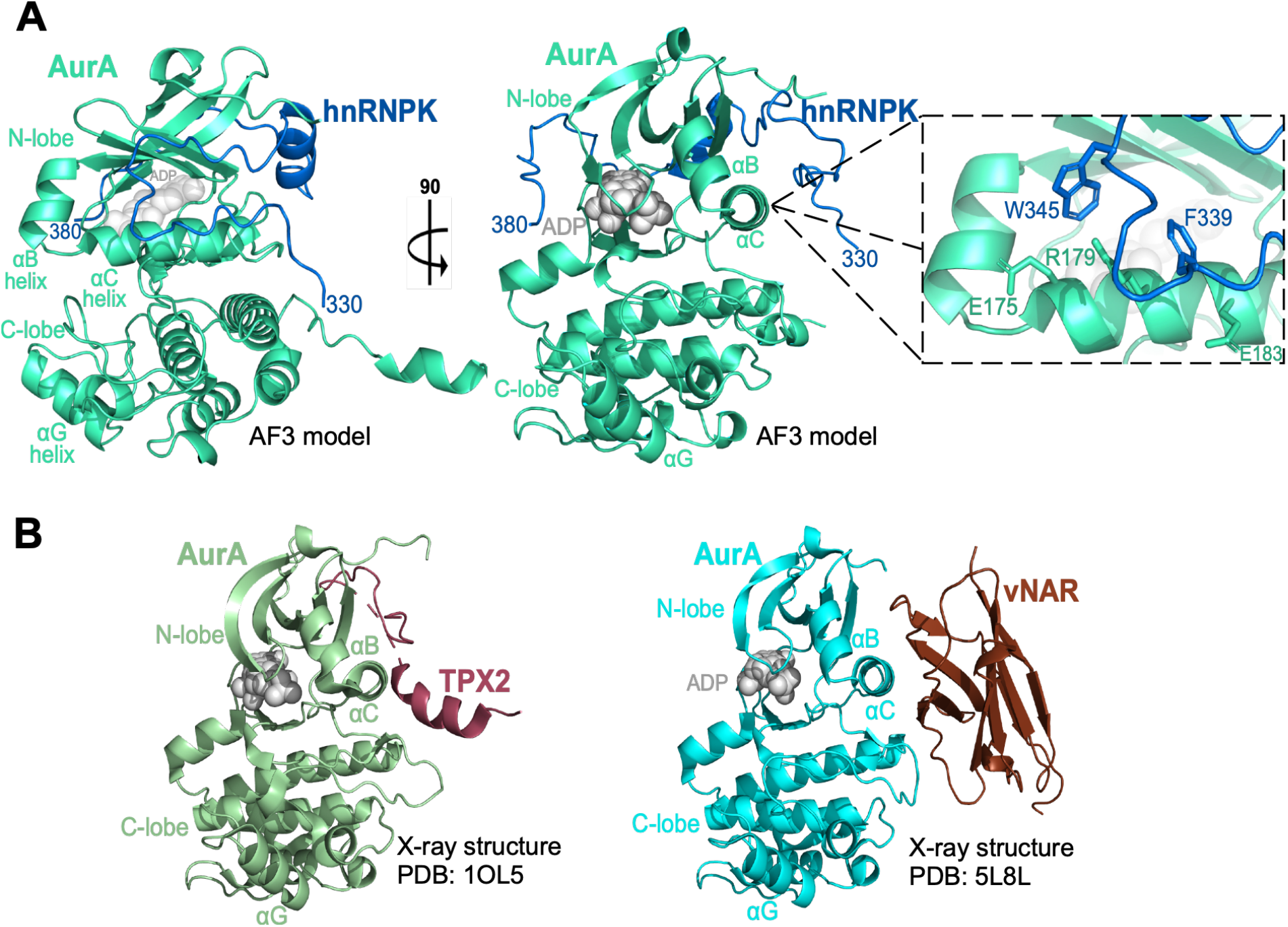
AlphaFold3 model of AurA in complex with hnRNPK^330-380^. **(A)** Cartoon representation of the AlphaFold3 model in different orientations with a magnified view of the key interaction site. **(B)** Structures of AurA in complex with vNAR-D01 (PDB 5L8L) and TPX2 (1OL5), in the same orientation representing overlapping binding sites.

As the binding site for hnRNPK on AurA in the AlphaFold3 model overlaps with the known binding site of TPX2, we used co-precipitation experiments to test whether these proteins exhibit competitive binding to AurA (Figure 3). His-TPX2^1-43^ and hnRNPK^1-463^ both independently co-precipitated with immobilised AurA, but when these partners were combined, the amount of hnRNPK^1-463^ that was co-precipitated was decreased in a dose-dependent manner by addition of His-TPX2^1-43^ (Figure 3A). Further competitive co-precipitation experiments were carried out using the AurA-binding regions of two other well-characterised binding-partners: transforming acidic coiled-coil protein 3 (TACC3) ^519-563^, which binds a separate site on the AurA N-lobe adjacent to the TPX2-binding site, and CEP192^468-533^, which binds across both these sites (Burgess et al., 2018; Holder et al., 2024). CEP192^468-533^ was observed to compete efficiently with hnRNPK, while His-TACC3^519-563^ did not affect the binding of hnRNPK to AurA (Figure S2A, S2B). These results are consistent with the observed competition between TPX2 and hnRNPK for binding to AurA and together the data confirm that, while the TACC3 pocket is not involved, the hnRNPK binding site does overlap with that of TPX2. This surface of the AurA N-lobe incorporates three ‘hotspots’ where aromatic side chains of TPX2 bind, termed the Y, F and W pockets (McIntyre et al., 2017). CEP192 interacts with the F and Y pockets, and also the TACC3 pocket (Burgess et al., 2018; Holder et al., 2024). Of these sites, hnRNPK is predicted by AlphaFold3 to bind at the Y-pocket, adjacent to the αC-helix, that is also recognised by Tyr8 and Tyr10 of TPX2 (Figure 3A). To test this prediction more precisely, competitive co-precipitation experiments were carried out using the AurA-directed single-domain antibody vNAR-D01, which binds only at the Y-pocket (Figure 2B) (Burgess et al., 2016). Co-precipitation of hnRNPK^1-463^ with AurA was decreased in a dose-dependent manner by addition of vNAR-D01, indicating that the Y-pocket is a major determinant of hnRNPK binding (Figure 3B).

**Figure 3.**
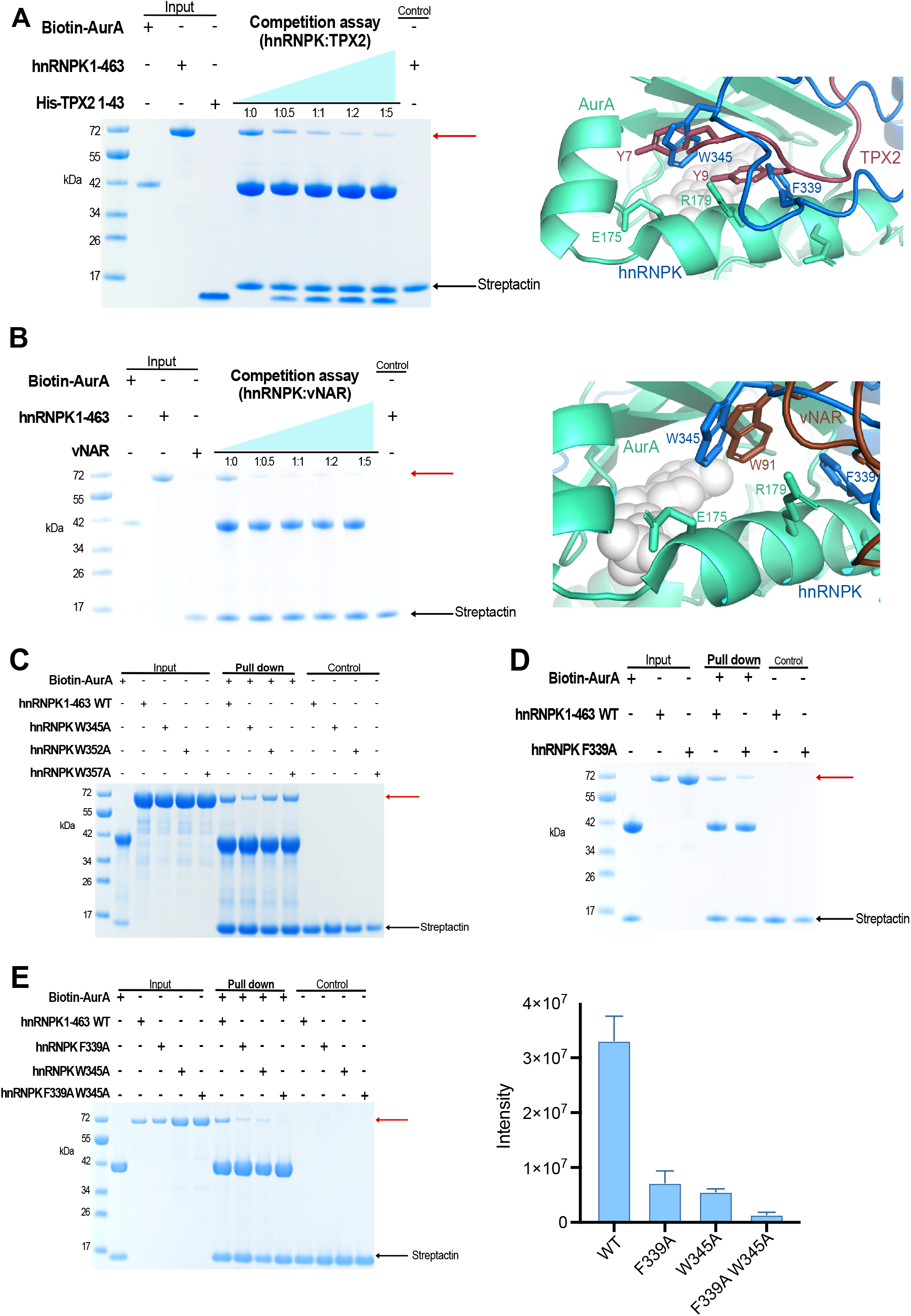
TPX2 and vNAR compete with hnRNPK for binding to AurA. Co-precipitation assays with biotinylated Avi-tagged AurA^122-403^, hnRNPK^1-463^, and His-TPX2^1-43^ **(A)** and vNAR-D01 **(B)**. 200 μg of Avi-AurA^122-403^ was immobilised on Streptactin Sepharose beads and incubated with 4 μM hnRNPK^1-463^ and 0, 2, 4, 8, and 20 μM His-TPX2^1-43^ or vNAR-D01. SDS-PAGE gels were used to visualise co-precipitated proteins (left). Bands corresponding to pulled-down hnRNPK are indicated by a red arrow. Structures of AurA in complex with vNAR-D01 (PDB 5L8L), and TPX2 (1OL5) are shown in the same orientation, representing overlapping binding sites (right). **(C)** Co-precipitation assays with biotinylated Avi-AurA^122-403^ and hnRNPK^1-463^ WT, and mutants (as indicated). Avi-AurA^122-403^ was immobilised on Streptactin Sepharose beads and incubated with hnRNPK proteins. SDS-PAGE gels were used to visualise co-precipitated proteins. Bands corresponding to pulled-down hnRNPK are indicated by a red arrow. **(D)** Co-precipitation assays with biotinylated Avi-AurA^122-403^ and hnRNPK^1-463^ WT and F339A mutant. **(E)** Co-precipitation assays with biotinylated Avi-AurA^122-403^ and hnRNPK^1-463^ WT and mutants (as indicated) (left). Histogram indicating quantification of hnRNPK protein co-precipitation by iBright analysis of SDS-PAGE gel band intensities, (n=3) error bars indicate SD (right).

The binding of vNAR-D01 to AurA is dependent on a single tryptophan residue, Trp91 which binds into the Y-pocket (Figure 3B), and its mutation to alanine was found to completely abolish binding (Burgess et al., 2016). There are three tryptophan residues in the amino acid sequence of hnRNPK, clustered in the disordered region between KH2 and KH3, Trp345, Trp352, and Trp357. These residues were each mutated to alanine and co-precipitation assays carried out to determine if these mutations disrupted the binding to AurA (Figure 3C). The hnRNPK W352A and W357A mutants exhibited no difference in AurA-binding compared to WT protein, while binding of the hnRNPK W345A mutant was significantly reduced (Figure 3C). This finding is consistent with the AurA titration NMR experiments which show strongest peak shifts and intensity losses for Trp345 (Figure 1D-F). These data indicate that Trp345 of hnRNPK is a key residue in the interaction with AurA and may bind into the Y-pocket in a similar manner to Trp91 of vNAR-D01, as predicted by the AlphaFold3 model (Figure 3B). Mutation to alanine of the other aromatic residue that the model predicts to be directly involved in the interaction, Phe339, also significantly reduced binding to AurA (Figure 3D) and binding of a F339A, W345A double mutant was almost completely eliminated (Figure 3E). These biochemical results together demonstrate that the interaction of hnRNPK with AurA is mediated by residues Phe339 and Trp345 of hnRNPK and the Y-pocket on the AurA N-lobe, which is consistent with the AlphaFold3 model presented.

As an AurA substrate protein that also interacts stably with AurA, hnRNPK has similarities to TACC3, the binding of which stimulates AurA catalytic activity in order to facilitate its own phosphorylation at a site adjacent to the AurA binding site (Burgess et al., 2015; Burgess et al., 2018). This example led us to test whether hnRNPK affects AurA catalytic activity. We tested phosphorylation of a Kemptide substrate by the AurA kinase domain in the presence of full-length hnRNPK or the 330-463 fragment in *in vitro* kinase assays (Figure 4A, Figure S3A). hnRNPK efficiently stimulated AurA-dependent substrate phosphorylation in both these experiments. Activatory EC50 values of 4.7 ± 0.4 μM and 2.0 ± 1.2 μM were measured for the full-length hnRNPK and hnRNPK^330-463^ proteins, respectively. As expected, hnRNPK mutants previously identified as having reduced AurA binding exhibited reduced stimulation of AurA kinase activity (Figure 4A). hnRNPK-mediated stimulation of AurA autoactivation was assayed by use of immunoblotting to monitor activation loop phosphorylation at Thr288 of initially unphosphorylated AurA. Briefly, increasing stoichiometric ratios of full-length hnRNPK (Figure 4B) or hnRNPK^330-463^ (Figure S3B) to dephosphorylated AurA, up to a ratio of 20 μM: 2.5 μM, were included in kinase assay reactions which were then analysed by SDS-PAGE and α-AurA pThr288 immunoblotting, and the intensities of bands were quantified and plotted. Full-length hnRNPK exhibited dose-dependent stimulation of AurA autophosphorylation with increasing stoichiometries up to 1:1, and dose-dependent inhibition of autophosphorylation at superstoichiometric concentrations (Figure 4B). hnRNPK^330-463^ also clearly stimulates AurA autophosphorylation, but dose-dependent effects were not observed (Figure S3B). The biphasic response to full-length hnRNPK may suggest that an N-terminal region of the protein has weak inhibitory activity.

**Figure 4.**
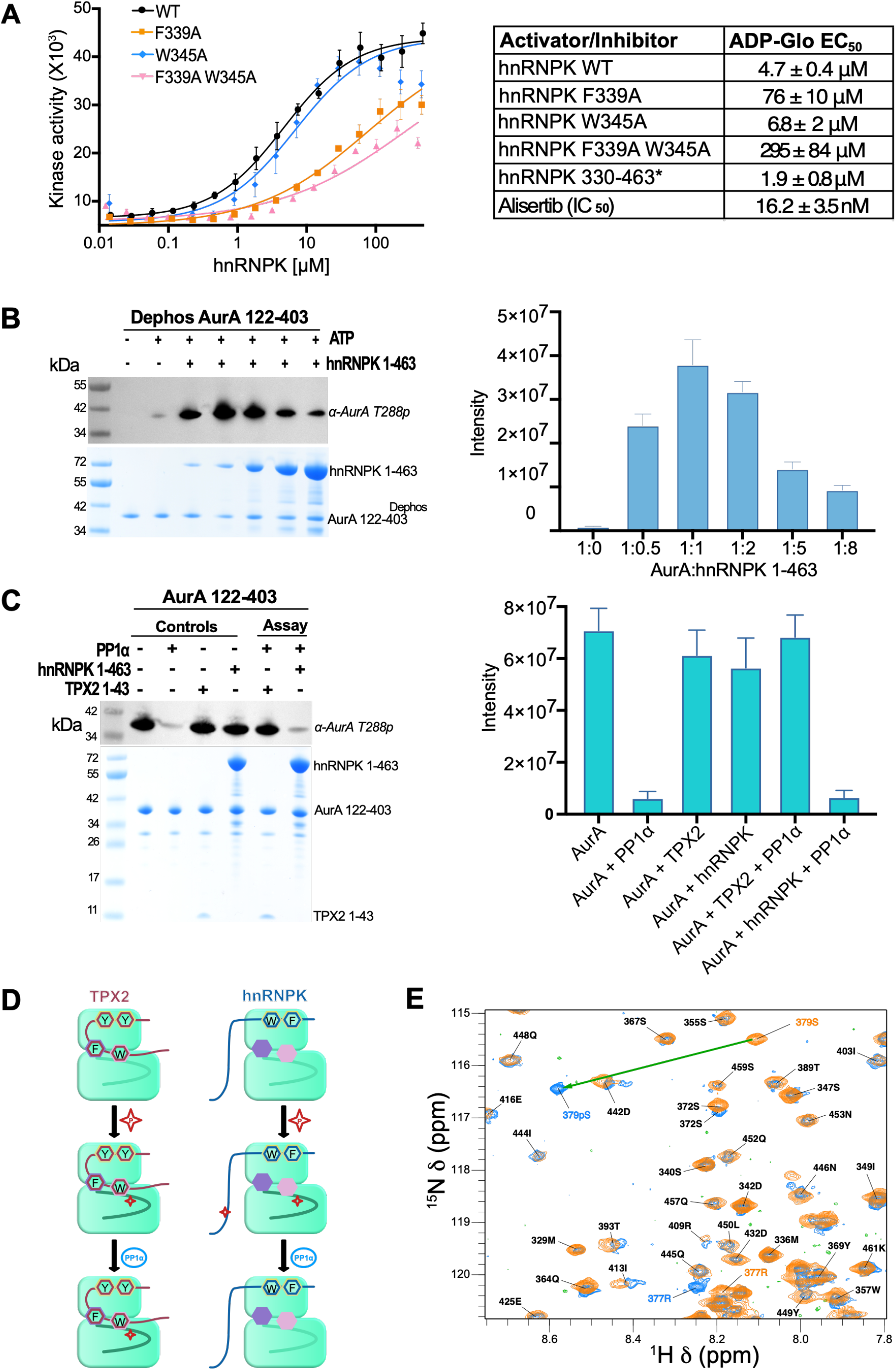
hnRNPK enhances AurA autophosphorylation and substrate phosphorylation. **(A)** Kinase assays to measure effect of hnRNPK on AurA substrate phosphorylation. 10 nM dephosphorylated AurA^122-403^ was incubated with 10 μM ATP, 100 μM of Kemptide and increasing concentrations of hnRNPK^1-463^ WT and mutants, as indicated. Alisertib, an AurA inhibitor, was used as a negative control. Assays were resolved using ADP-Glo kit (Promega). Luminescence data points represent the average of three experiments, and error bars indicate SD. **(B)** Assays to measure effect of hnRNPK on AurA autophosphorylation. 2.5 μM of unphosphorylated AurA^122-403^ was incubated with 160 μM ATP and 0, 1.25, 2.5, 5, 12.5 and 20 μM of hnRNPK^1-463^. Assays were visualised SDS-PAGE and levels of phosphorylation were observed by immunoblotting using an α-AurA pThr288 antibody (Cell Signalling Technology) (left). Histogram displaying quantification of AurA phosphorylation at Thr288 quantified from band intensity by iBright analysis (n=3) (right). Error bars indicate SD. **(C)** Phosphatase protection assay. 3 μM phosphorylated AurA^122-403^ was incubated with 0.1 μM phosphatase PP1α in the absence or presence of 10 μM hnRNPK^1-463^. The assay was visualised by SDS-PAGE and AurA Thr288 phosphorylation observed by immunoblotting using an α-AurA pThr288 antibody (left). Histogram displaying quantification of AurA phosphorylation at Thr288 quantified from band intensity by iBright analysis (n=3) (right). Error bars indicate SD. **(D)** Schematic showing difference in protection/phosphorylation of TPX2 compared to hnRNPK. Binding pockets on AurA are shown as coloured hexagons; Y (yellow), F (purple) and W (pink) pockets **(E)** Phosphorylation of hnRNPK Ser379 by AurA shown by changes in ^1^H–^15^N HSQC spectra: a large downfield shift in the Ser379 peak and small local chemical shift perturbations.

As association with TPX2 protects the AurA activation loop from dephosphorylation by phosphatases (Bayliss et al., 2004), a phosphatase protection assay was carried out to investigate whether hnRNPK carries similar properties (Figure 4C). Phosphorylated AurA was incubated with full-length hnRNPK and the phosphatase PP1α, which dephosphorylates AurA at the pThr288 site. AurA Thr288 phosphorylation was monitored by immunoblotting and band intensities quantified. While the presence of TPX2^1-43^ preserved pThr288 as expected, Thr288 phosphorylation was significantly decreased by PP1α both in the absence and presence of hnRNPK, indicating that hnRNPK does not protect AurA from dephosphorylation. This observation is consistent with the AlphaFold3 model of the complex as TPX2-mediated protection of AurA from dephosphorylation is known to require regions of TPX2 that bind at the F- and W-pockets (Bayliss et al., 2004) while in the model, hnRNPK is bound at the Y-pocket only (Figure 4D).

NMR experiments were used to track the phosphorylation of hnRNPK^330-463^ using a catalytic amount of WT AurA. Ser379 was phosphorylated by AurA, confirming the observation made by others (Hsueh et al., 2011; Tsai et al., 2019) (Figure 4E). Phosphorylation was fast and specific to Ser379. Cα shift measurements indicated that there was no change in structure upon phosphorylation except for Ser379 and surrounding residues (Figure S4), and hnRNPK peaks showed a similar profile of intensity loss upon addition of AurA, indicating that AurA-binding is unaffected by phosphorylation.

## Discussion

Previously published studies identified hnRNPK as a substrate phosphorylated by AurA on Ser379 and their interaction was suggested to be part of a mechanism by which AurA promotes cancer cell migration (Hsueh et al., 2011; Tsai et al., 2019). AurA and hnRNPK were also shown to cooperate in transcriptional activation of *MYC*, a gene that is overexpressed in a majority of human cancers (Naso et al., 2021). In the present work we have characterised the structural basis and biochemical consequences of the interaction.

hnRNPK is an activator of AurA kinase activity, enhancing both autoactivation and substrate phosphorylation, and this interaction is mediated by binding of a short intrinsically disordered region of hnRNPK, adjacent to the Ser379 phosphorylation site, at a pocket on the N-lobe of AurA that has previously been identified as a part of the TPX2 binding site termed the Y-pocket. Residues Phe339 and Trp345 of hnRNPK are key residues in the interaction, and modelling studies suggest that their aromatic side chains mimic the binding mode of those of Tyr8 and Tyr10 of TPX2, or Trp91 of the AurA-directed single-domain antibody vNAR-D01, within this pocket. Unlike TPX2, hnRNPK does not protect the AurA activation loop from dephosphorylation by phosphatases because it binds at the Y-pocket only, while regions that bind at the F- and W-pockets are required for TPX2 to protect pThr288. The AlphaFold3 model presented here is consistent with all experimental data and thus serves as a plausible representation for the physiological complex.

This work provides tools that could be applied to explore the function of the AurA/hnRNPK complex in cancer cell models. For example, the hnRNPK F339A, W345A double mutant that we have identified, which cannot bind AurA, could now be exploited to identify phenotypes associated with targeted loss of its interaction with AurA. Similarly, as we have demonstrated that vNAR-D01, an inhibitor of the AurA-TPX2 interaction, also disrupts the AurA-hnRNPK interaction, it follows that tools developed to target the AurA-TPX2 interaction could be employed to inhibit the AurA-hnRNPK interaction in cancer cell models. An abundance of nuclear AurA is hallmark of cancer, but AurA lacks an NLS (Naso et al., 2021). Therefore, it is likely that the interactions of AurA with nuclear binding partners such as hnRNPK and N-Myc is what drives nuclear localisation of AurA in cancer cells (Richards et al., 2016). Furthermore, formation of the AurA/hnRNPK complex on chromatin has been shown to allow these proteins to cooperate to direct oncogenic transcriptional activity (Zheng et al., 2016). Exclusion of AurA from the nucleus or suppression of aberrant gene expression might be accomplished by disruption of this complex, and the advancements in structural and molecular understanding of the interaction outlined here may facilitate future studies aimed at targeting it for therapeutic purposes.

## Methods

### Cloning, recombinant protein expression and purification

The vectors pET30TEV AurA^122-403^ C290A C393A, pGEX-cs TPX2^1-43^, pET30TEV TPX2^1-43^, pETM6T1 C-terminally tagged Avi-AurA ^122-403^ were produced in earlier work (Bayliss et al., 2003; Rowan et al., 2013; Burgess et al., 2015; Burgess et al., 2016). AurA^122-403^ C290A C393A, Avi-AurA, AurA^122-403^ WT and GST-TPX2^1-43^ were expressed and purified as stated in earlier work (Bayliss et al., 2003; Burgess et al., 2015; Burgess et al., 2016).

Expression vectors for hnRNPK constructs were transformed into BL21 (DE3) *E. coli* cells and grown in LB media at 37 °C until an induction OD600 of approximately 0.6 was obtained and 0.5 mM IPTG added. Cultures were incubated overnight at 21°C prior to cell harvesting by centrifugation. Protein purification was performed using a HisTrap HP column as per the manufacturer’s instructions. The His-tag was removed by overnight TEV cleavage. A final SEC step was performed with protein loaded onto a HiLoad 16/600 Superdex 200 pg column (Cytiva) equilibrated in 150 mM NaCl, 20 mM Tris pH 7.5, 2 mM BME and 10% glycerol.

For ^15^N/^13^C- or ^15^N-labelled samples of hnRNPK^330-463^, LB cultures (1 L) were grown to an OD600 ∼0.6 as described above, but cells were then pelleted (2600*g*, 25 min), resuspended/washed in PBS buffer to remove residual LB, pelleted again and resuspended in 250 ml of ^15^N/^13^C- or ^15^N-labelled minimal media. Following a 1-h incubation period at 21 °C, overnight expression, cell harvesting and protein purification were carried out as described above for unlabelled protein. In the final SEC step, an alternative NMR buffer was used (20 mM K2HPO4/KH2PO4 pH 6.5, 150 mM NaCl, 2 mM BME). Minimal media contained 2 g/L ^15^NH4Cl and 6–8 g/L ^13^C or unlabelled D-glucose in 50 mM Na2HPO4, 25 mM KH2PO4, 20 mM NaCl, supplemented with 2 mM MgSO4, 0.2 mM CaCl2, 0.01 mM FeSO4, a micronutrient cocktail and a vitamin solution (‘BME’ 100× solution, Sigma–Aldrich).

### Size-exclusion chromatography

AurA was applied to a Superose 12 10/300 GL column alone or with an interacting protein. The complexes were made in a 1:1.2 molar ratio of Aurora-A to interacting protein and incubated on ice for 1 h prior to chromatography. Fractions were visualised using SDS-PAGE analysis.

### Co-precipitation assays

200 μg of biotinylated Avi-AurA^122-403^ was incubated with 25 μL of Strep-Tactin Sepharose resin (Cytiva) in 500 μL of binding buffer for 30 min. Beads were washed with 5 mL binding buffer and incubated with 200 μg of interacting protein for 1 h at 4 °C with agitation. The resin was washed with 5 mL binding buffer. Bound proteins were analysed by SDS-PAGE and visualised by Coomassie Blue staining.

### Competition co-precipitation assays

200 μg of biotinylated Avi-AurA^122-403^ was incubated with 25 μL of Strep-Tactin Sepharose resin (Cytiva) in 500 μL of binding buffer (50 mM Tris pH 7.5, 300 mM NaCl, 5 mM β-mercaptoethanol, 5 mM MgCl2, 10 % glycerol and 0.1 % Tween 20) for 30 min. Beads were washed with 5 mL binding buffer and incubated with 4 μM of hnRNPK and 0, 2, 4, 8, or 20 μM of CEP192^468-533^, His-TACC3^519-563^, vNAR-D01 or His-TPX2^1-43^ for 1 h at 4 °C on a roller. The resin was washed with 5 mL binding buffer. Bound proteins were analysed by SDS-PAGE and visualised by Coomassie Blue staining and by immunoblotting using anti-his antibody (Sigma-Aldrich).

### Kinase assay

An ADP-Glo (Zegzouti et al., 2009) kit was used to measure the activity of AurA substrate phosphorylation upon binding to hnRNPK. 500 μM hnRNPK^1-463^ or hnRNPK^330-463^ were serially diluted across 16 wells, and each dilution included in a kinase assay mix also containing 10 nM dephosphorylated AurA^122-403^, 10 μM ATP, and 100 μM Kemptide (Cambridge Bioscience Ltd) that was incubated at room temperature for 1 h in white 384-well low volume low binding plates (Greiner Bio-one). 5 μL of each assay was incubated with 5 μL of ADP-Glo reagent for 40 min at room temperature to remove excess ATP and stop the kinase reaction. 10 μL of Kinase detection reagent was added and incubated for 30 min at room temperature, to convert ADP to ATP. The ATP is then quantified by luminescence from an ATP-dependent luciferase reaction. Luminescence was recorded using an HIDEX plate reader with a 1 second integration time. Data points represent the average of three experiments, and error bars indicate SD. Graphs were fitted using the log(agonist) vs. response – (three parameters) equation in Prism10 (GraphPad) to calculate EC50 values reflecting kinase activation. Results are reported as EC50 ± SE. All ADP-Glo™ reagents were purchased from Promega.

### Autophosphorylation assay

2.5 μM AurA^122-403^, co-expressed with λ phosphatase and purified to homogeneity, was incubated with 160 μM ATP, alone and in the presence of 0, 1.25, 2.5, 5, 12.5 and 20 μM of interacting protein for 1 h at room temperature. Assays were carried out in 20 mM Tris pH 7.5, 25 mM NaCl, 1 mM MgCl2, 1 mM DTT and 0.01 % Tween 20. Reactions were stopped by the addition of 2X SDS-loading buffer and separated by SDS-PAGE. Immunoblots were generated and probed with an α-AurA pThr288 antibody (Cell Signalling Technology).

### Phosphatase protection assay

3 μM AurA^122-403^, expressed in the absence of λ phosphatase, was incubated with 0.1 μM PP1α alone and in the presence of 10 μM of hnRNPK^1-463^ for 1 h at room temperature. Assays were carried out in 50 mM Tris pH 7.5, 0.1 mM EDTA, 2 mM MgCl2, 5 mM DTT and 0.025% Tween 20. Reactions were stopped by the addition of 2X SDS-loading buffer and separated by SDS-PAGE. Immunoblots were generated and probed with an α-AurA pThr288 antibody (Cell Signalling Technology).

### AlphaFold3 modelling

AlphaFold3 (Abramson et al., 2024) was used through the AlphaFold server for models of hnRNPK^330-380^ in complex with AurA. Automated seed function was used, with 20 iterations each generating 5 models, to output a total of 100 models. The alignment function in PyMOL was used to compare all models. Of these models, 28 models with iPTM scores ranging from 0.77 to 0.8 were similar. Structure figures were generated using PyMOL (The PyMOL Molecular Graphics System, Version 2.3.5 Schrödinger, LLC).

### NMR spectroscopy

NMR spectra were recorded with a 300 µM sample of ^15^N/^13^C-labelled hnRNPK 330-463 in NMR buffer containing 5% D2O and a protease inhibitor cocktail solution using a 600-MHz Oxford Instruments magnet equipped with a Bruker Avance console, a QCI-P-cryoprobe and Topspin software. ^1^H–^15^N HSQC spectra were recorded at 10, 15, 20, 25 and 30 °C. Peak intensities were generally improved at 25 °C compared to 10 °C. For assignment, HNCO, HNcaCO, HNCA, HNCoCA HNCACB and ^1^H–^13^C HSQC spectra were recorded at 25 and 10 °C (all using 20% NUS). Additional HNcocaCB and HBHAcoNH spectra were recorded at 10 °C. Data were processed with NMRPipe (Delaglio et al., 1995), assignments and further analysis used CCPNmr Analysis (v2.5) (Vranken et al., 2005). Spectra allowed *ab initio* assignment of the ^1^H–^15^N HSQC spectrum and 96% of possible HN, H, CO, Cα, and C β resonances at 25 °C (77% were recorded at 10 °C). Assignments in the KH3 domain were validated by comparison with BMRB entry 4405 (a hnRNPK KH3 domain mutant). Chemical shift data were deposited in the Biological Magnetic Resonance Data Bank (BMRB) entry 53201. A structural analysis using all measured chemical shifts was carried out using the TALOS-N web server (https://spin.niddk.nih.gov/bax-apps/nmrserver/talosn/ (Shen and Bax, 2013).

**For AurA titrations**, small aliquots of concentrated (600 µM) AurA 122-403 C290A, C393A were added to ^15^N- or ^15^N/^13^C-labelled samples of hnRNPK 330-463 (50–100 µM) and changes tracked within ^1^H–^15^N HSQC or ^1^H–^15^N TROSY spectra. Chemical shift perturbations (CSPs) were calculated using a 0.15 weighting for ^15^N shifts: CSP^2^ = ΔH^2^ + (0.15 × ΔN)^2^.

**For *in situ* phosphorylation**, a 90 µM sample of ^15^N/^13^C-labelled hnRNPK 330-463 in NMR buffer, supplemented with 1 mM ATP and 2.5 mM MgCl2, was treated with 2 µM AurA 122-403 WT. ^1^H–^15^N HSQC spectra were recorded before and after addition. Complete conversion, as gauged by loss of the Ser379 peak and accumulation of a pSer379 peak, was achieved within the time taken to record a spectrum (<40 min). To confirm assignment of pSer379 and locally shifted peaks in the ^1^H–^15^N HSQC spectrum, and to generate Cα assignments, a HNCA experiment was recorded. For structural comparison after phosphorylation, Cα secondary shifts (Δδ = δ – δref) were calculated using reference coil values (δref) available online (https://www1.bio.ku.dk/english/research/bms/sbinlab/randomchemicalshifts2/ (Hendus-Altenburger et al., 2019).

## Supplementary

**Figure S1.**
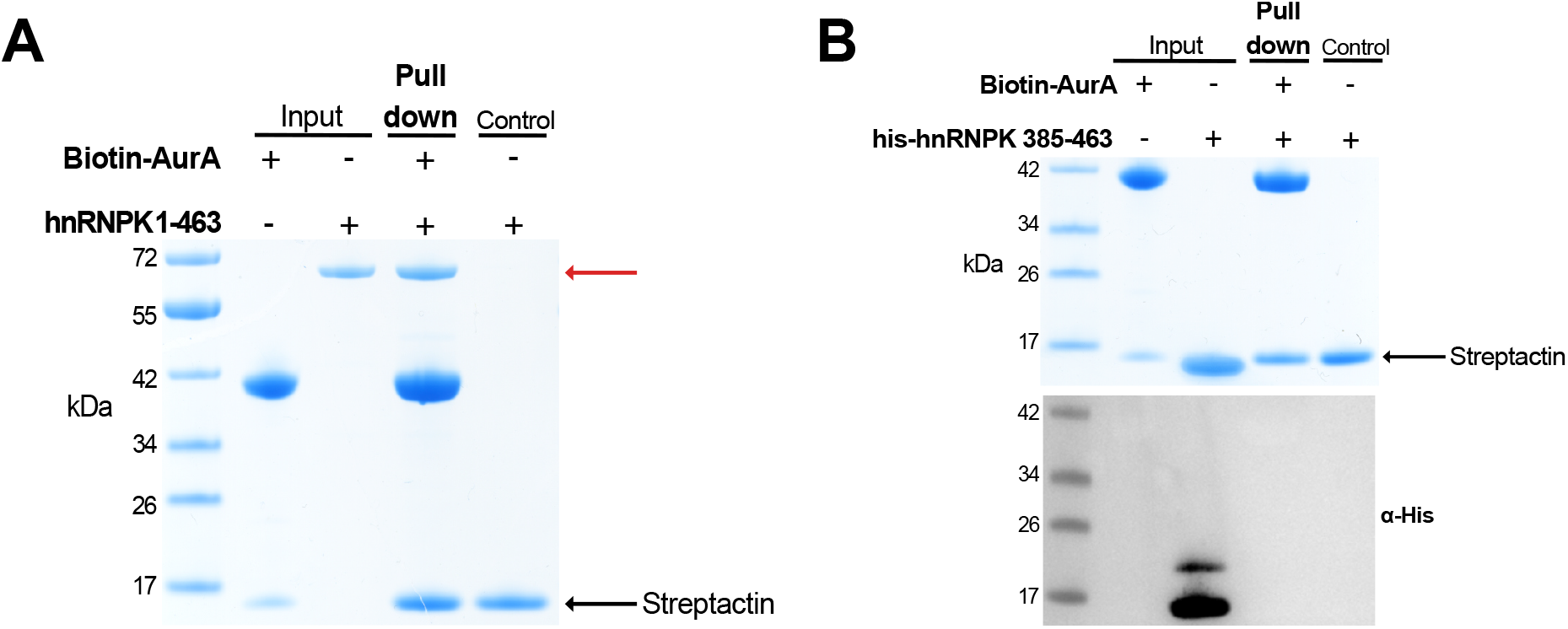
Full length hnRNPK, but not the C-terminal KH3 domain, binds to AurA. Co-precipitation assay with biotinylated Avi-tagged AurA^122-403^ and hnRNPK^1-463^ **(A)** and His-tagged hnRNPK^385-463^ KH3 domain **(B)**. Avi-AurA^122-403^ was immobilised on Streptactin Sepharose beads and incubated with hnRNPK proteins. SDS-PAGE gels were used to visualise co-precipitated protein. Bands corresponding to hnRNPK are indicated by a red arrow. The shorter hnRNPK^385-463^ KH3 domain protein was visualised by α-His immunoblotting because it migrates to a similar degree as streptactin.

**Figure S2.**
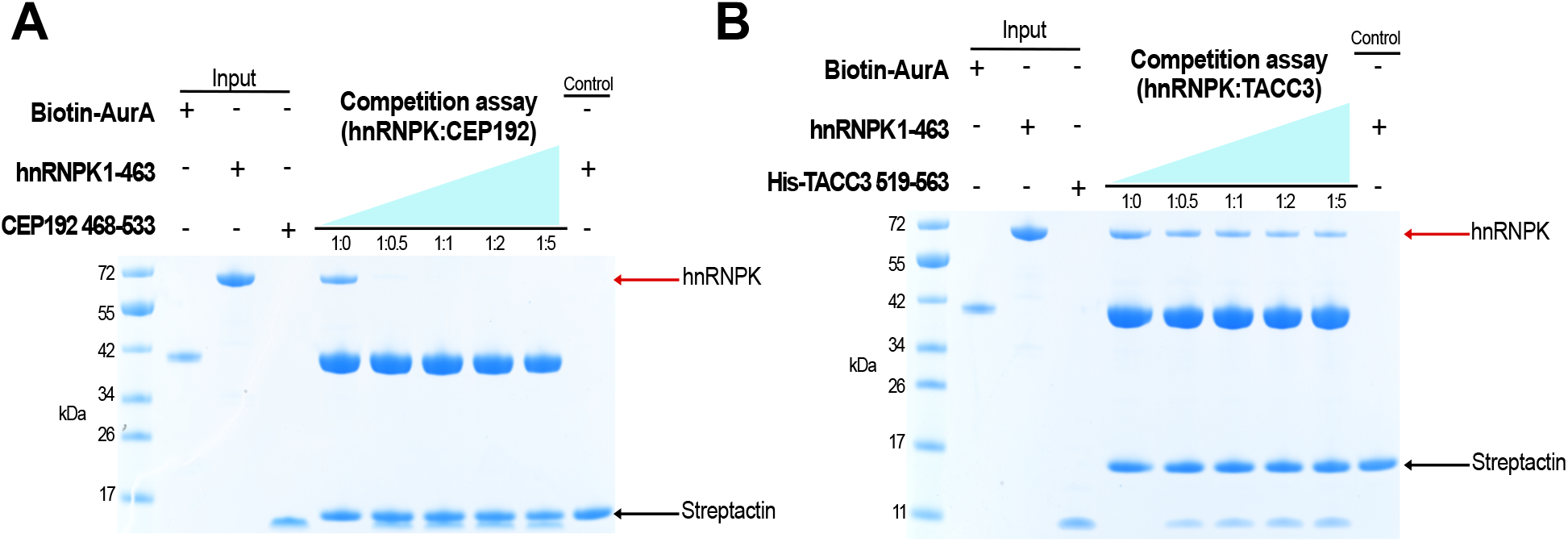
Co-precipitation assay with CEP192 and TACC3. Co-precipitation assays using immobilised biotinylated Avi-AurA^122-403^ and hnRNPK^1-463^ with CEP192^468-533^**(A)** or with His-TACC3^519-563^**(B)**. 200 μg of Avi-AurA^122-403^ was immobilised on Streptactin Sepharose beads and incubated with 4 μM hnRNPK and 0, 2, 4, 8, or 20 μM of CEP192^468-533^ or His-TACC3^519-563^. SDS-PAGE gels were used to visualise co-precipitated protein.

**Figure S3.**
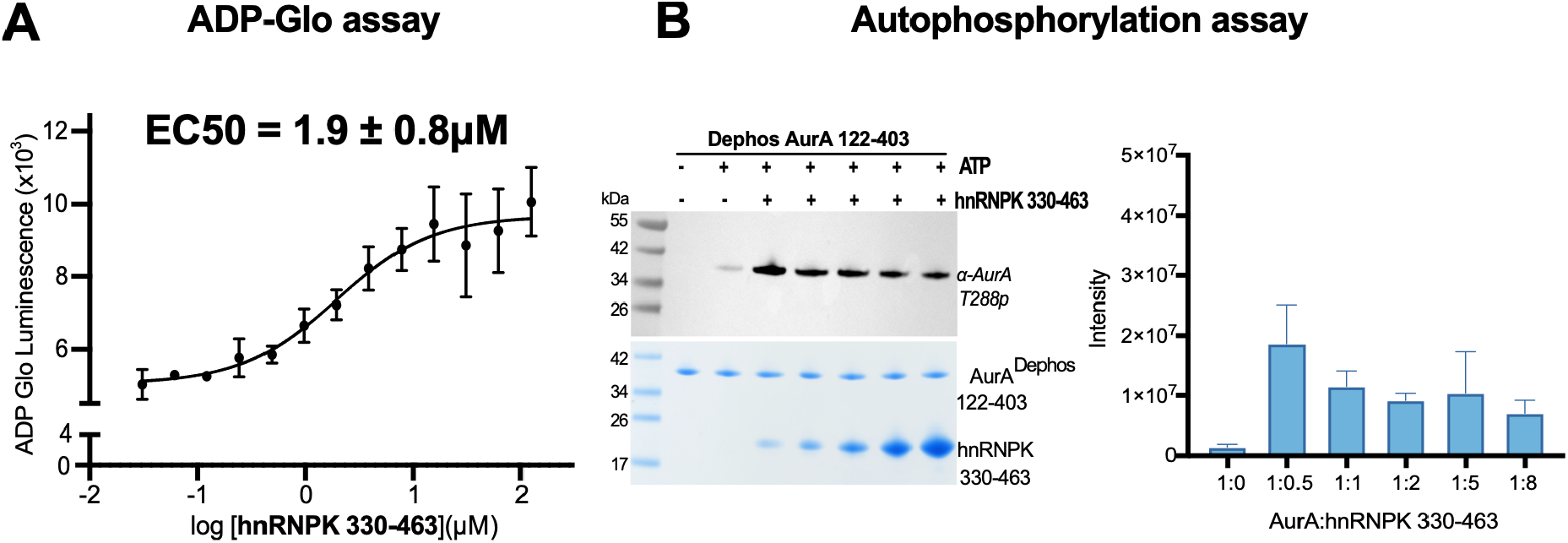
The C-terminal 330-463 residues of hnRNPK enhance the autophosphorylation of AurA. **(A)** 10 nM phosphorylated AurA^122-403^ was incubated with 10 μM ATP, 100 μM of kemptide and increasing concentrations of hnRNPK^330-463^. Assays were resolved using ADP-Glo kit (Promega). Luminescence data points represent the average of three experiments, and error bars indicate SD. Graphs were fitted using the log(agonist) vs. response – (three parameters) equation in Prism10 (GraphPad) to calculate EC50 values reflecting kinase activation. EC50 value 2.0 ± 1.2 μM were measured. **(B)** Autophosphorylation assay in which 2.5 μM of unphosphorylated AurA^122-403^ was incubated with 160 μM ATP and 0, 1.25, 2.5, 5, 12.5 and 20 μM of hnRNPK^330-463^. Assays were visualised with SDS-PAGE gels and levels of phosphorylation were observed by immunoblotting, using an α-AurA pThr288 antibody (CST). Plot displaying quantification of AurA phosphorylation at Thr288. Bands were quantified using iBright analysis (n=3). Error bars indicate SD.

**Figure S4.**
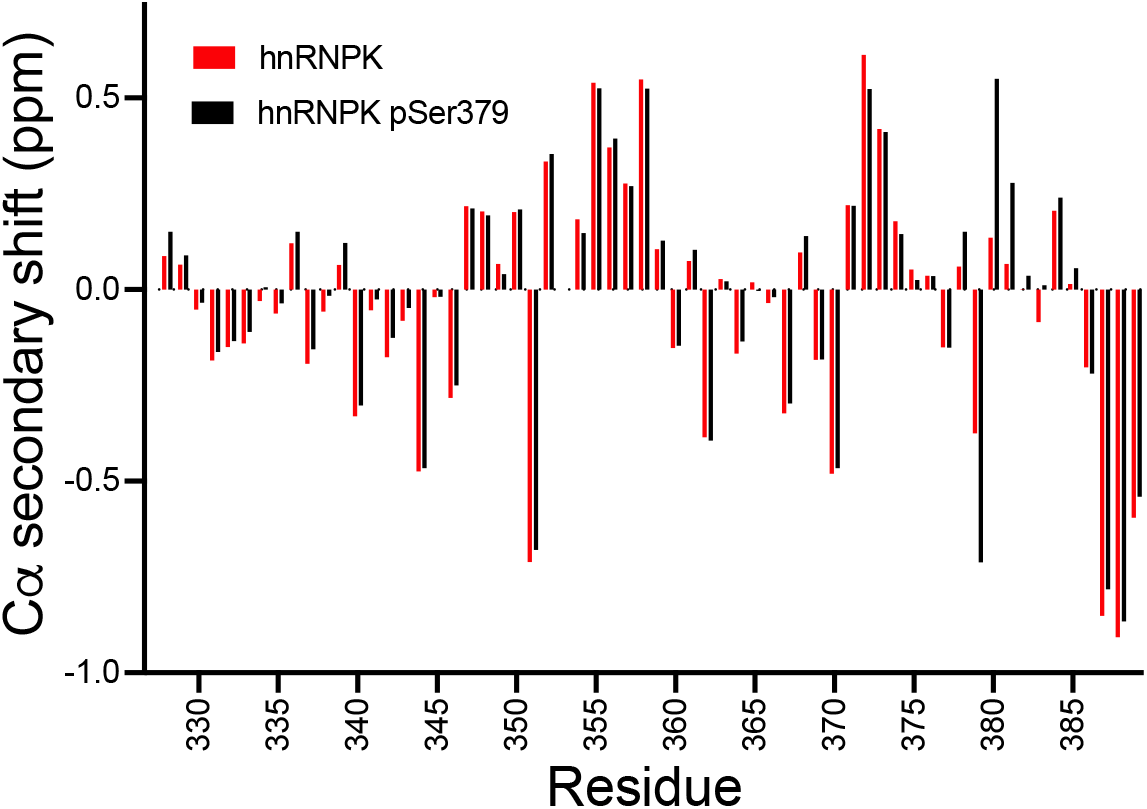
The effect of Ser379 phosphorylation on hnRNPK structure. Secondary shifts for Cα nuclei were measured before (red) and after phosphorylation of Ser379 with AurA (black).

## Author contributions

**IAM:** Investigation, Conceptualisation, Methodology, Formal analysis, Writing - Original draft, Visualisation, Writing – review and editing. **LMV:** Investigation. **MB:** Investigation, Formal analysis, Writing – review and editing. **MWR:** Supervision, Conceptualisation, Validation, Visualisation, Formal analysis, Writing - Original draft, Writing – review and editing. **TO:** Supervision. **RB:** Conceptualisation, Supervision, Writing – review and editing, Funding Acquisition.

## Acknowledgments

We thank Dr Selena Burgess for providing the His-TACC3^519-563^, WT AurA^122-403^ and vNAR-D01 proteins, and Dr Jennifer Miles for providing the CEP192^468-533^ proteins used in this work. We thank Arnout Kalverda for NMR support and Roland Dunbrack Jr (Fox Chase Cancer Centre) for help with modelling. Facilities for NMR spectroscopy were funded by the University of Leeds (ABSL award) and Wellcome Trust (108466/Z/15/Z). We acknowledge funding from BBSRC (BB/V003577/1, BB/V003577/2) and Cancer Research UK (C24461/A23302).

